# Chicken cGAS senses fowlpox virus infection and regulates macrophage effector functions

**DOI:** 10.1101/2020.10.01.321422

**Authors:** Marisa Oliveira, Damaris Ribeiro Rodrigues, Vanaique Guillory, Emmanuel Kut, Efstathios S. Giotis, Michael A. Skinner, Rodrigo Guabiraba, Clare E Bryant, Brian J Ferguson

## Abstract

The anti-viral immune response is dependent on the ability of infected cells to sense foreign nucleic acids. In multiple species, the pattern recognition receptor (PRR) cyclic GMP-AMP synthase (cGAS) senses viral DNA as an essential component of the innate response. cGAS initiates a range of signalling outputs that are dependent on generation of the second messenger cGAMP that binds to the adaptor protein stimulator of interferon genes (STING). Here we show that in chicken macrophages, the cGAS/STING pathway is essential not only for the production of type-I interferons in response to intracellular DNA stimulation, but also for regulation of macrophage effector functions including the expression of MHC-II and co-stimulatory molecules. In the context of fowlpox, an avian DNA virus infection, the cGAS/STING pathway was found to be responsible for type-I interferon production and MHC-II transcription. The sensing of fowlpox virus DNA is therefore essential for mounting an anti-viral response in chicken cells and for regulation of a specific set of macrophage effector functions.

## Introduction

The ability of virally infected cells to mount an effective innate immune response is dependent on the intracellular sensing of nucleic acids by pattern recognition receptors (Mansur et al., 2014). The PRRs that sense and respond to intracellular DNA are well characterised in a number of mammalian and non-mammalian organisms but are less studied in avian species, including chickens (Bryant et al., 2015). The PRR cyclic cAMP-GMP (cGAMP) synthase (cGAS) binds intracellular viral DNA and, via production of the second-messenger 2’3’-cGAMP, triggers a range of signalling outputs including type-I interferon (IFN-I) production, cell death and cellular senescence (Li and Chen, 2018). The absence of cGAS or the adaptor protein, stimulator of interferon genes (STING), which binds cGAMP, results in the susceptibility to DNA virus infection in knockout mice and impairs IFN-I production by cells infected with DNA viruses or transfected with linear double stranded DNA (Li et al., 2013). Through its ability to sense mislocalised self-DNA, the cGAS/STING signalling axis is also a potent regulator of autoinflammatory and anti-tumour immune responses (Ablasser et al., 2013a; Mullard, 2017). People with activating mutations in STING or loss-of function mutations in the 5’-3’ exonuclease TREX, which removes excess cytoplasmic dsDNA, suffer from interferonopathies (Crow and Rehwinkel, 2009).

The ability of cGAS/STING signalling to drive multiple downstream signalling outputs is dependent on the activation of a number of distinct signalling mechanisms, some of which are better defined than others. The production of IFN-I in this context is dependent on STING recruiting and facilitating activation of TANK-binding kinase-1 (TBK1) and the transcription factor interferon regulatory factor-3 (IRF3) (Tanaka and Chen, 2012). IRF3 phosphorylation, dimerisation and translocation to the nucleus results in IFN-I transcription. The mechanism or mechanisms by which STING can promote cell death are less well described, but include inflammasome activation (Gaidt et al., 2017) and apoptosis of various cell types including myeloid and T cells (Gulen et al., 2017; Sze et al., 2013). cGAS can also activate a programme of cellular senescence in fibroblasts by sensing damaged self-DNA (Glück et al., 2017). It is not currently clear in what contexts these disparate signalling outputs are activated by cGAS/STING and to what extent they cross-talk with each other.

Chickens are economically important livestock birds that are infected by numerous viruses including fowlpox virus (FWPV). Fowlpox is a virus from the *poxviridae* family that replicates its double stranded DNA genome in the cytoplasm of infected cells. The infection is characterised by proliferative lesions in the skin that progress to thick scabs (cutaneous form) and by lesions in the upper GI and respiratory tracts (diphtheritic form) (Giotis and Skinner, 2019). Transmitted mechanically by biting insects, it causes significant losses to all forms of poultry production systems (from backyard, through extensive to intensive commercial flocks). It is particularly challenging in tropical climes where control of biting insects is difficult. FWPV is also used as a live recombinant vaccine vector in avian and mammalian species (Lousberg et al., 2010). Like other poxviruses the cytoplasmic replication cycle of FWPV exposes large amounts of foreign DNA to intracellular DNA sensing PRRs, making cGAS a likely candidate for sensing FWPV infection and making FWPV a potentially useful tool for delineating nucleic acid sensing mechanisms in avian systems. The mechanisms by which FWPV is sensed by PRRs during infection have not, however, been described.

In this study we show the existence of a cGAS/STING pathway in chicken macrophages and determine its downstream signalling outputs. Using cGAS and STING CRISPR/Cas9 knockout HD11 cells and pharmacological inhibitors of STING and TBK1 in primary macrophages, we show that the activation of cGAS by intracellular DNA drives a IFN-I response and that this response can be enhanced by priming cells with IFNα. As well as driving IFN-I production, we show that cGAS/STING signalling in macrophages can enhance transcription of specific immune recognition molecules including genes encoding the class II major histocompatibility complex (MHC-II) and co-stimulatory proteins, but without altering phagocytosis. Using FWPV mutants that are deficient in specific immunomodulators we are able to overcome the immunosuppression of wild type FWPV and show that this virus is sensed by cGAS, resulting in IFN-I and MHC-II transcription. These data show that the cGAS/STING/TBK1 pathway senses viral DNA in chicken macrophages and that this pathway regulates not only the antiviral interferon response but also modulates specific components of macrophage effector function machinery.

## Materials and Methods

### Reagents

Calf Thymus (CT) DNA (Sigma), Herring Testes (HT) DNA (Sigma), polyinosinic-polycytidylic acid (poly(I:C), Invivogen), 2’3’-cGAMP (Invivogen) and chicken interferon alpha (Yeast-derived Recombinant Protein, Kingfisher Biotech, Inc) were diluted in nuclease-free water (Ambion, ThermoFisher). H-151 and BX795 (Invivogen) were diluted in DMSO, following the manufacturer’s protocols.

### Cell Culture

HD11 cells, an avian myelocytomatosis virus (MC29)-transformed chicken macrophage-like cell line (Beug et al., 1979), were incubated at 37°C, 5% CO_2_. They were grown in RPMI (Sigma-Aldrich, Germany) complemented with 2.5% volume per volume (v/v) heat-inactivated foetal bovine serum (FBS; Sera Laboratories International Ltd), 2.5% volume per volume (v/v) chicken serum (New Zealand origin, Gibco, Thermo Fisher Scientific), 10% Tryptose Phosphate Broth solution (Gibco, Thermo Fisher Scientific), 2 mM L-glutamine (Gibco, Thermo Fisher Scientific), 50 µg/mL of penicillin/streptomycin (P/S; Gibco, Thermo Fisher Scientific).

Chicken embryonic fibroblasts (CEFs) (Pirbright Institute, Woking, UK) were incubated at 37°C, 5% CO_2_ and were grown in Dulbecco’s Modified Eagle Medium (DMEM) -F12 with Glutamax (Gibco), 5% v/v FBS, and 50 µg/mL P/S.

### Knock-out HD11 cell line generation by CRISPR-Cas9

#### CRISPR guide design

According to the *MB21D1* (cGAS) and *TMEM137* (STING) sequences obtained from the Ensembl database (release 94), single guide (sg)RNA sequences (Table 1) were designed targeting the catalytic domain (residues 11-13 and 109) and start of the open reading frame, for cGAS and STING, respectively.

**Table 1.**
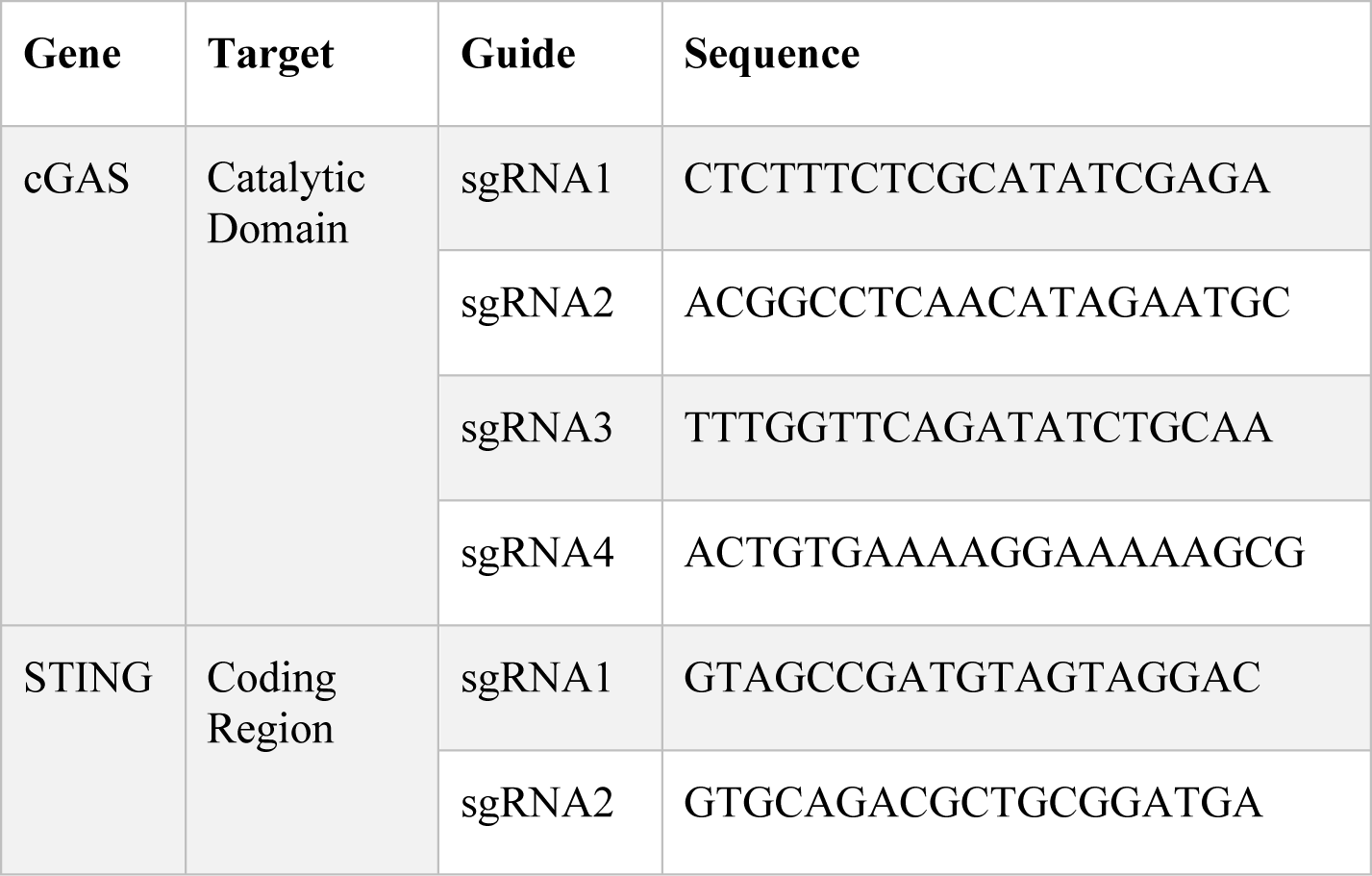
CRISPR/Cas9 guide RNAs

### Knock-out cell lines generation using CRISPR-Cas9

Genome editing of HD11 was performed using ribonucleoprotein (RNP) delivery. tracrRNA was mixed with the target specific sgRNA (Table 1), followed by an incubation at 95°C. To form the RNP complex, the tracrRNA/sgRNA mix was incubated with the Cas9 protein (IDT, Leuven, Belgium) and electroporation enhancer at 21°C.

To generate knockout cells, 1×10^6^ cells per guide were electroporated with the corresponding RNP complex using Lonza Electroporation Kit V (Lonza). After 48 h, the cells were expanded for future experiments and their DNA were extracted using the PureLink Genomic DNA Kit (Thermo Scientific, Waltham, MA, USA). The knockout efficiency was evaluated by genotyping the polyclonal cell populations using MiSeq (Illumina) according to a published method (Schmidt et al., 2016). The primers used for the sequencing are listed Table 2.

**Table 2.**
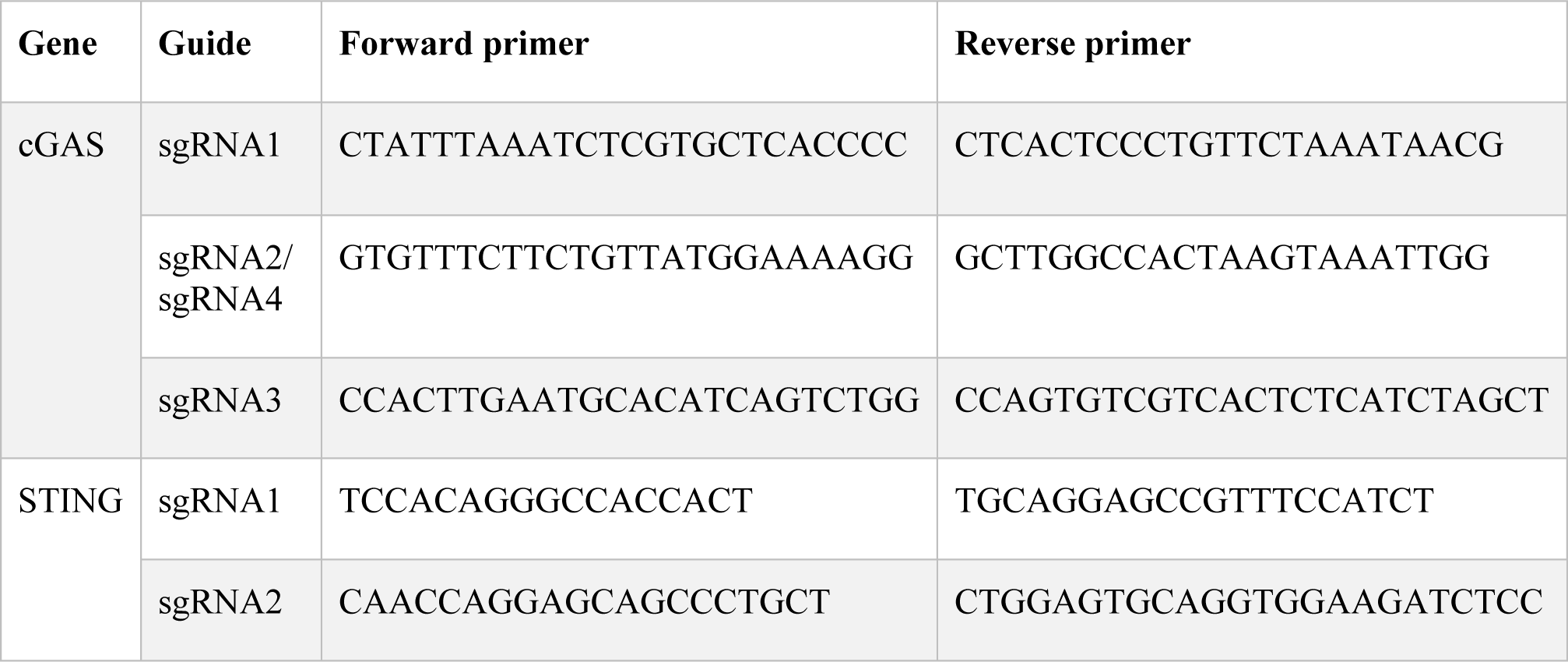
Illumina sequencing primers

The successfully edited populations (using guides cGAS sg3 and STING sg1) were diluted to a concentration of 0.5 cell/well and seeded in 96-well plates. Individual clones were sequenced by MiSeq and the confirmed knockout clones were expanded for experiments.

### Primary macrophages

Chicken bone marrow derived macrophages (BMDM) were generated as previously described (Garrido et al., 2018). Briefly, femurs and tibias of 4 week-old immunologically mature White Leghorn (PA12 line) outbred chickens were removed, both ends of the bones were cut and the bone marrow was flushed with RPMI supplemented with P/S. Cells were then washed and re-suspended in RPMI, loaded onto an equal volume of Histopaque-1077 (Sigma-Aldrich, Germany), and centrifuged at 400 g for 20 min. Cells at the interface were collected and washed twice in RPMI. Purified cells were seeded at 1 × 10^6^ cells/ml in sterile 60 mm bacteriological petri dishes in RPMI supplemented with 10% FBS, 25 mM HEPES, 2 mM L-glutamine, P/S and 25 ng/ml recombinant chicken colony stimulating factor 1 (CSF-1) (Kingfisher Biotech, Inc) at 41 °C and 5% CO2. Half of the medium was replaced with fresh medium containing CSF-1 at day 3. At day 6, adherent cells were harvested and cultured in RPMI supplemented with 10% FBS, 25 mM HEPES, 2 mM L-glutamine, and P/S prior to stimulation.

### Stimulation Assays

HD11 (WT, cGAS and STING knockouts) were seeded in 12-well plates at a density of 3 × 10^5^ cells/well. In the following day, the cells were transfected using TransIT-LT1 (Mirus Bio, USA) with HT-DNA (1, 2 or 5 μg/mL), CT-DNA (1, 2 or 5 μg/mL) or Poly(I:C) (1 μg/mL), and harvested 6 h or 16 h post-transfection. In the priming assays, IFNα (200 ng/mL) was added 16 h hours prior to transfection. 2’3’ cGAMP was added at a concentration of 2.5 μg/mL and cells were harvested 6 h post-treatment.

BMDM were seeded in 6-well plates at 8×10^5^ cells/ml. In the following day, cells were transfected using TransIT-LT1 with HT-DNA (2 μg/mL), CT-DNA (2 μg/mL) or Poly(I:C) (1 μg/mL), and harvested 6 h post-transfection. In the priming assays, IFNα (50 ng/ml) was added 16 h prior transfection to the cells supernatants. 2’3’ cGAMP was added to cells supernatants at the concentration of 10 μg/mL and the cells were harvested 6 h post-treatment.

### Chicken IFN-I bioassay

The presence of IFN-I in supernatants of stimulated BMDM was measured indirectly using a luciferase-based Mx-reporter bioassay (Schwarz et al., 2004). Briefly, cells from the quail fibroblast cell line CEC32 carrying the luciferase gene under the control of chicken Mx promoter (kindly provided by Prof. Peter Stäheli, University of Freiburg, Germany) were seeded at 2.5 × 10^5^ cells/well in 24-well plates and incubated at 41 °C under 5% CO_2_. The next day, cells were incubated for 6 h with the diluted supernatants (1/10 of total volume). Medium was removed and cells were washed twice with PBS. Cells were lysed using the Cell Culture Lysis Reagent (Promega, USA), according to the manufacturer’s instructions, and luciferase activity was measured using the Luciferase assay reagent (Promega, USA) and a GloMax-Multi Detection System (Promega, USA).

### Cell viability

BMDM viability following different stimuli was assessed using the fluorescent DNA intercalator 7-aminoactinomycin D (7-AAD, BD Biosciences, USA). Briefly, following stimulations, supernatants were discarded and the cells were harvested and washed in PBS. Cells were stained according to the manufacturer’s protocol and the viability was analyzed by flow cytometry (BD FACS Calibur). Data were expressed as the percentage of 7AAD positive cells over total acquired events (50,000 cells).

### RNA Extraction

Cells were lysed by overlaying with 250 µL of lysis buffer containing 4 M guanidine thiocyanate, 25 mM Tris pH 7, and 143 mM 2-mercaptoethanol. As a second step, 250 µL of ethanol was added, and the solution was transferred to a silica column (Epoch Life Science, Inc., Sugar Land, TX, USA) and centrifuged; all centrifugation steps were performed for 90 seconds at 16600 *g*. The bound RNA was washed by centrifugation with 500 µL of buffer containing 1 M guanidine thiocyanate, 25 mM Tris pH 7, and 10% ethanol, followed by a double washing step with 500 µL of wash buffer 2 (25 mM Tris pH 7 and 70% (v/v) ethanol). RNA was eluted by centrifugation in 30 µL of nuclease-free water and the concentration was measured using a NanoDrop 2000 Spectrophotometer (Thermo Scientific, Waltham, MA, USA).

### cDNA and qPCR

Using 500 ng of RNA extracted from HD11 cells, cDNA was produced using SuperScript III reverse transcriptase, following the manufacture’s protocol (Thermo Scientific, Waltham, MA, USA). Samples were diluted in nuclease-free water in a 1:2.5 ratio. 1 μl of the diluted product was used for quantitative PCR (qPCR) in a final volume of 10 μl. qPCR was performed using SybrGreen Hi-Rox (PCR Biosystems Inc.) using primers described in Table 3. Fold change in mRNA expression was calculated by relative quantification using hypoxanthine phosphoribosyltransferase (HPRT) as endogenous control.

**Table 3.**
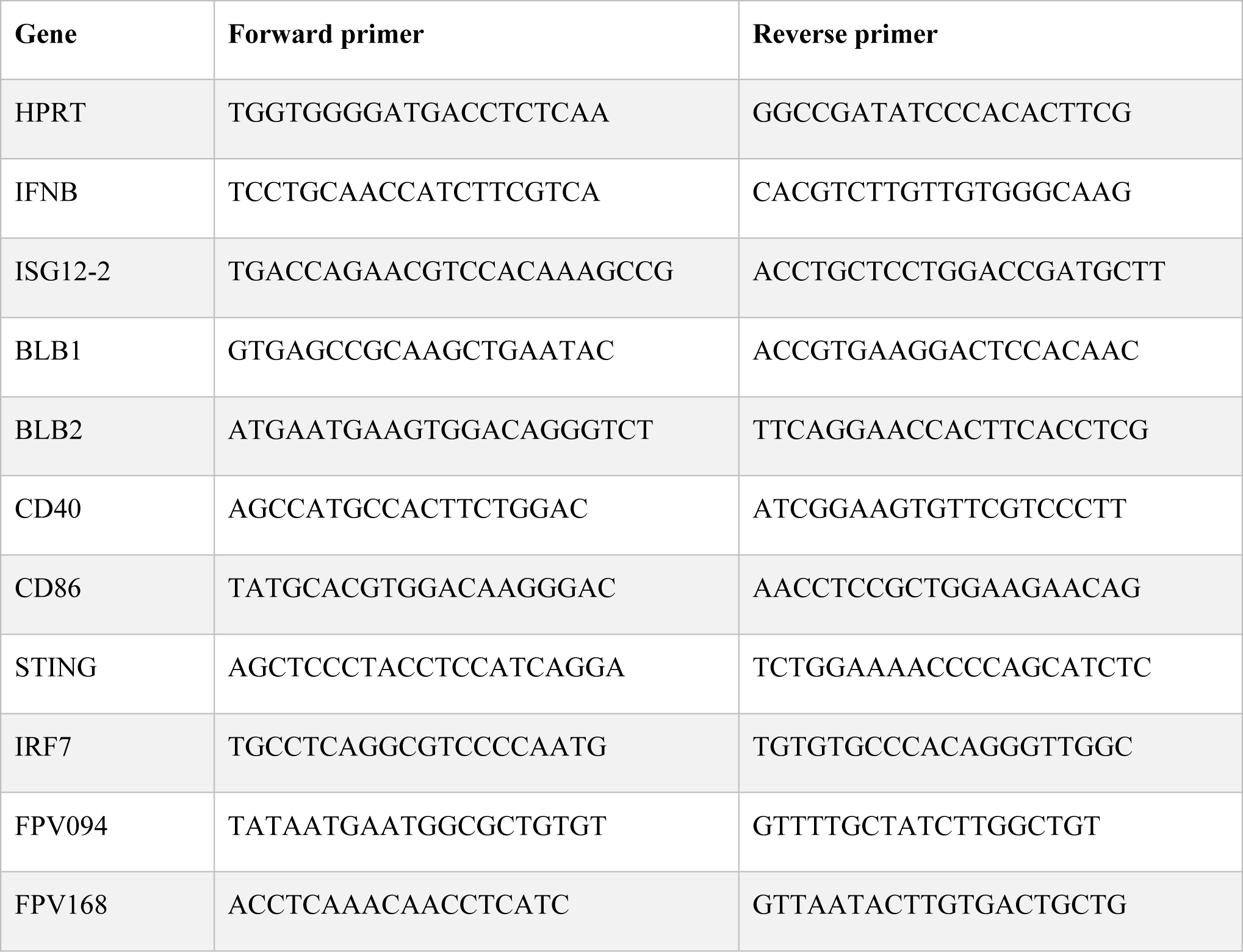
qRT-PCR primer sequences

Total RNA (up to 1 µg per reaction) from BMDM was reverse transcribed with iScript cDNA synthesis kit (Bio-Rad, USA). Quantitative PCR was performed using 1 µl of cDNA, 5 µl of iQ SYBR Green Supermix (Bio-Rad, USA), 0.25 µl of each primer pair and 3.5 µl of nuclease-free water in a total reaction volume of 10 µl. Fold-increase in gene expression was calculated by relative quantification using HPRT and Glyceraldehyde 3-phosphate dehydrogenase (GAPDH) as endogenous controls.

### Phagocytosis assay by flow cytometry

HD11 WT cells were seeded at a confluence of 3×10^5^ cells/ml in 12-well plates. The cells were primed with IFNα for 16h and then with transfected exogenous DNA (HT-and CT-DNA – 2 μg/mL) or treated with 2’3’cGAMP (5 μg/mL) for 6 h. After this, the cells were incubated with Zymosan coated beads conjugated with FITC at a ratio of 30 beads to 1 cell for all conditions for 40 min at 37 °C. The cells were wash two times in PBS and fixed in suspension using the solution (missing ref; BD Biosciences) with 4% PFA. Cell populations were counted by analysis on a CytoFLEX cytometer.

### Fowlpox virus growth and titration

Fowlpox WT (FP9) and mutants (FPV012 (Laidlaw et al., 2013) and FPV184 (Giotis & Skinner, unpublished)) were propagated in primary chicken embryonic fibroblasts (CEFs) and grown in DMEM-F12 (Thermo Fisher Scientific, Waltham, MA, USA) containing 1% FBS and 5% P/S, and harvested 5 days later. 10-fold dilutions of cell supernatants were prepared in serum-free DMEM-F12 and used to inoculate confluent monolayers of CEFs for 1.5 h at 37°C. Cells were then overlaid with 2xMEM:CMC (1/1 ratio). The foci were counted 7 days later after staining with Toluidine Blue.

### Fowlpox virus infection

HD11 cells were seeded in 12-well plates in the day prior infection. Fowlpox viruses were diluted in serum-free DMEM-F12 at a multiplicity of infection (MOI) of 3 and added in the cells (1 ml per well). Infected cells and supernatants were collected from infections at 8h and 24h post-infection.

### Statistical Analysis

Prism 7 (GraphPad) was used to generate graphs and perform statistical analysis. Data were analyzed using an unpaired t test with Welch’s correction unless stated otherwise. Data with P < 0.05 was considered significant and 2-tailed P-value were calculated and presented as: <0.0001 - ****, >0.0001 - ***, >0.001 - **, >0.01 - *. Each experiment has at least two biological replicates unless stated.

## Results

### Intracellular DNA activates a IFN-I response in chicken macrophages

In order to assess the ability of chicken macrophages to sense and respond to intracellular DNA we used a combination of the monocytic cell line HD11 and primary bone marrow derived macrophages (BMDM). Transfection of CT DNA, increasing doses of HT-DNA or the RNA analogue poly(I:C) into HD11 cells resulted in transcription of chicken interferon-β (IFNβ) and the interferon stimulated gene (ISG) ISG12.2, an orthologue of mammalian IFI6 (Figure 1A). A dose-dependent response to DNA was observed. Transfection of DNA into primary BMDMs also resulted in IFNβ and ISG12.2 transcription (Figure 1B) and IFN-I secretion as measured by a bioassay (Figure 1C), indicating that this response is present in both primary macrophages and the transformed monocytic HD11 cell line.

**Figure 1.**
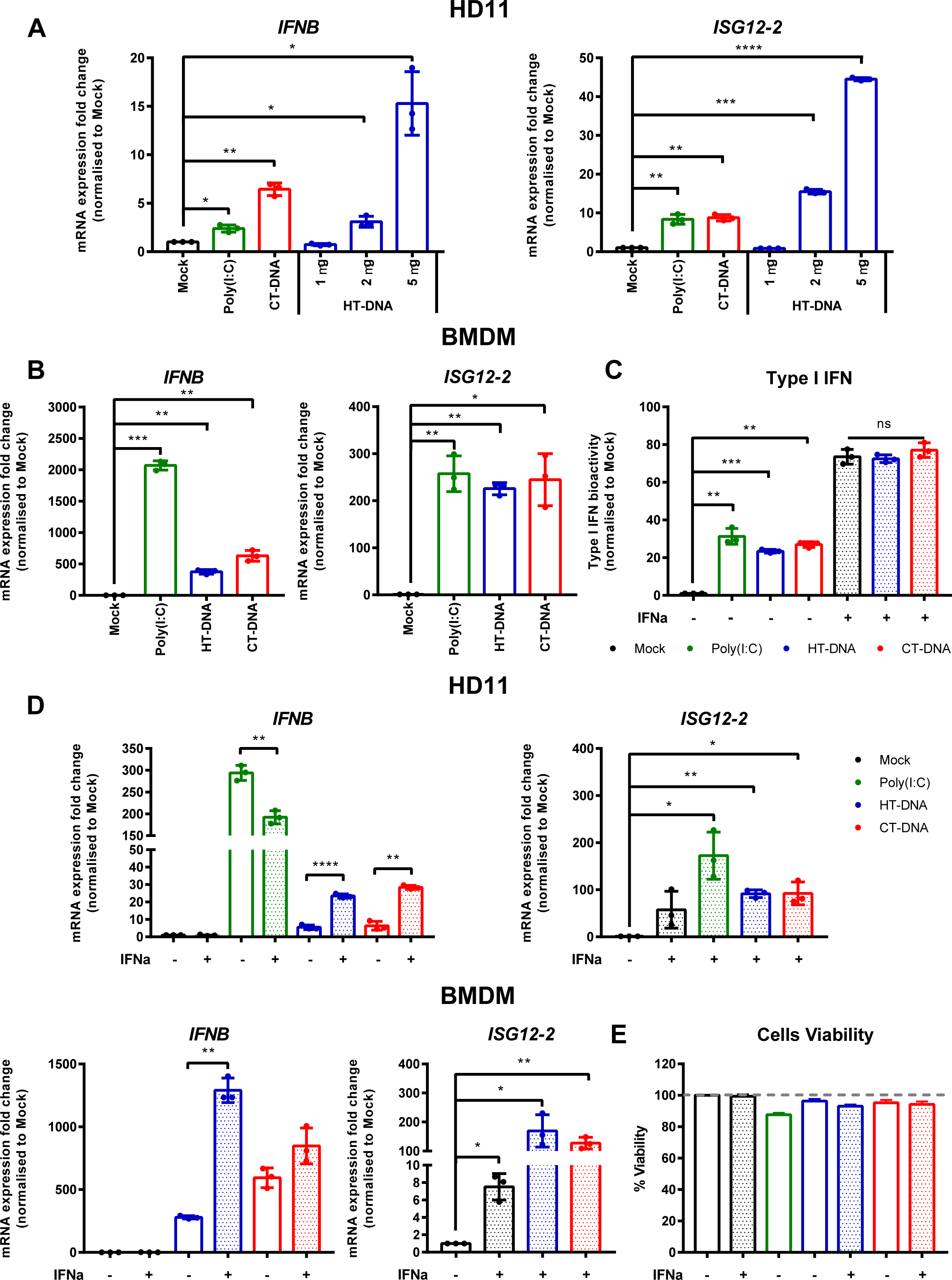
Intracellular DNA activates an IFN-I response in chicken macrophages. (A) HD11 cells were transfected with HT-DNA (1, 2 and 5 µg/mL), CT-DNA (5 µg/mL) or Poly(I:C) and transcription of IFNB and ISG12.2 measured by qRT-PCR 6 h later. (B) Chicken BMDM were transfected with HT-DNA, CT-DNA (2 µg/mL) or Poly(I:C) (1 µg/mL) and transcription of IFNB and ISG12.2 measured by qRT-PCR 6 h later. (C) Resting BMDMs or BMDMs primed with IFNα for 6 h were transfected with HT-DNA, CT-DNA (2 µg/mL) or Poly(I:C) (1 µg/mL) and interferon activity in the supernatants was measured after 24 h using a bioassay. (D) HD11 or BMDM were primed with IFNα for 6h, transfected with HT-DNA, CT-DNA, or Poly(I:C) and transcription of IFNB and ISG12.2 measured by qRT-PCR 6 h later. (E) BMDM were primed with IFNα for 6h, transfected with HT-DNA, CT-DNA, or Poly(I:C) and cell viability measured by 7AAD staining 24 h later. *: p < 0.05, **: p < 0.01, ***: p < 0.001; ****: p < 0.0001; ns: no significant difference.

Since in mammalian systems STING is recognised as an interferon stimulated gene (ISG) (Ma et al., 2015), we sought to understand the effect of IFN-I priming of macrophages on the response to intracellular DNA. Pre-treatment of HD11 or BMDM with chIFNα resulted in an enhancement of IFNβ transcription following DNA stimulation and confirmed ISG12.2 as an ISG (Figure 1D). This signalling enhancement might be explained by increased transcription of STING and/or IRF7 following IFNα treatment (Supplementary Figure 1). Across all HD11 and BMDM DNA stimulations we found that there was little observable or measurable cell death (Figure 1E), indicating that, in chicken macrophages, cell death is not a specific output of STING signalling.

### Intracellular DNA stimulates transcription of MHC-II and co-stimulatory molecules

The sensing of both intracellular and extracellular pathogens activates macrophages, causing up-regulation or enhancement of effector functions designed to combat infection. We hypothesised that DNA transfection, mimicking the presence of intracellular infection, might result in direct effects on the molecules that contribute to T cell stimulation. There are two chicken MHC-II beta chain genes, BLB1 and BLB2, both of which were transcriptionally upregulated by DNA stimulation in chicken BMDMs (Figure 2A). In HD11 cells, BLB1 transcription was upregulated by DNA stimulation, while BLB2 transcription was upregulated only by IFNα pre-treatment (not shown), highlighting possible differences between primary and transformed cells in this specific context (Figure 2B). CD86 and CD40 are key co-stimulatory molecules in T cell activation. In BMDM CD86 and CD40 transcription was upregulated in response to DNA stimulation (Figure 2A). There was, however, no measurable impact of DNA stimulation on phagocytosis as measured by bead-uptake assays in HD11 cells (Figure 2C). As such, key molecules involved in T cell activation by macrophages are regulated by DNA stimulation, but not all macrophage effector functions are equally enhanced by this signal.

**Figure 2.**
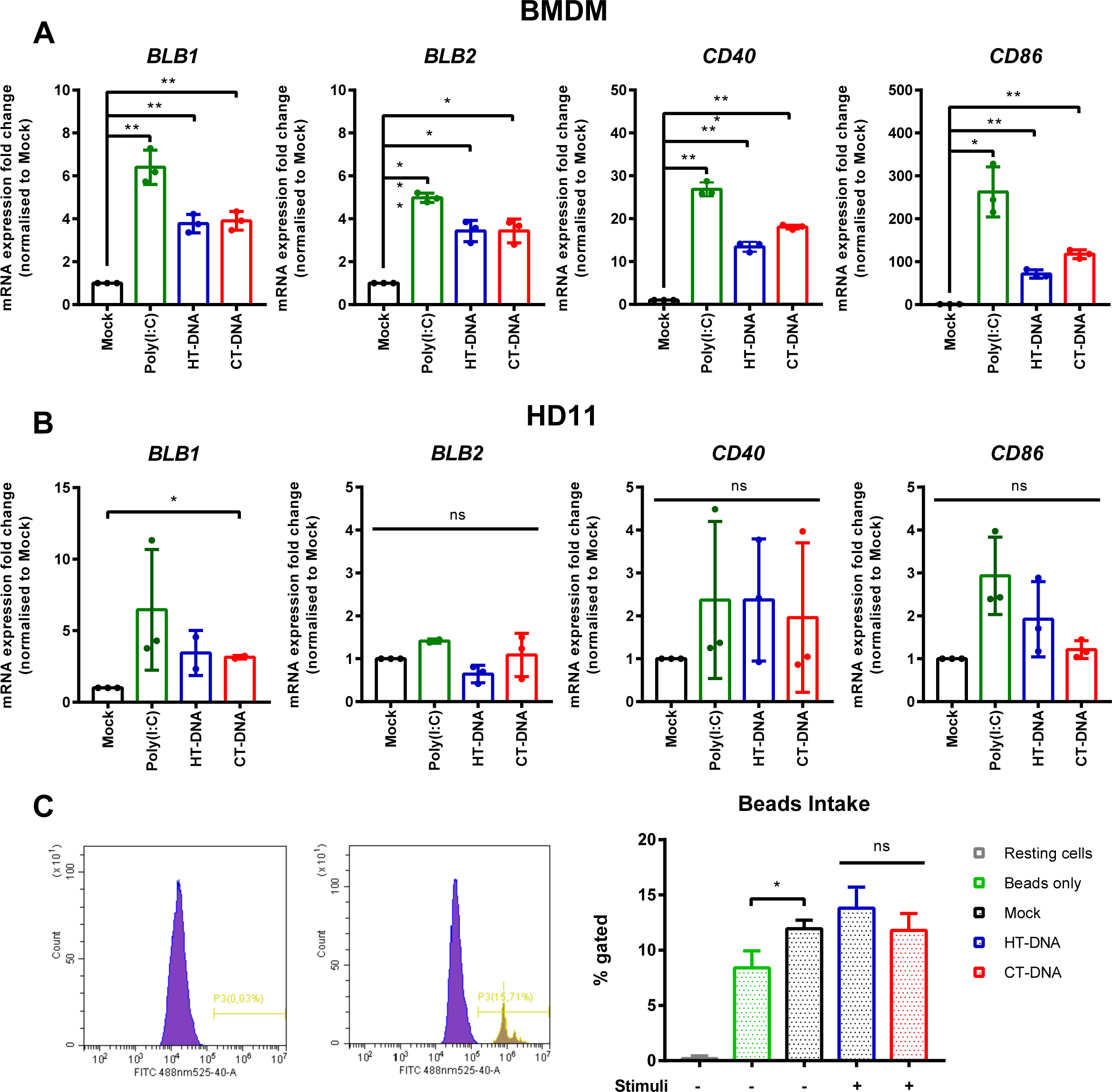
Intracellular DNA stimulates transcription of MHC-II and co-stimulatory molecules. (A) BMDMs or (B) HD11 cells were transfected with HT-DNA, CT-DNA, or Poly(I:C) and transcription of BLB1, BLB2, CD40 and CD86 measured by qRT-PCR 6 h later. (C) HD11 cells were stimulated with HT-DNA, CT-DNA, or Poly(I:C) and 6 h later phagocytosis was monitored by FITC-conjugated, zymosan coated bead uptake. Histograms of non-treated versus treated cells (left panels) and respective percentages of FITC positive cells for each treatment tested (right panel) are presented. *: p < 0.05, **: p < 0.01, ***: p < 0.001; ****: p < 0.0001; ns: no significant difference.

### STING and TBK1 contribute to DNA-driven transcriptional responses in chicken BMDMs

In order to dissect the signalling pathway downstream of intracellular DNA sensing, we first used the ligand 2’3’-cGAMP, the enzymatic product of cGAS that directly binds and activates STING (Ablasser et al., 2013b). Treatment of BMDMs or HD11 cells with 2’3’-cGAMP led to increased transcription levels of IFNb, ISG12.2, BLB1, BLB2, CD86 and CD40 (Figure 3A,B). This response, and the response to DNA stimulation, could be reduced by small molecule inhibitors of STING (H151) and the kinase TBK1 (BX795), indicating the existence of a STING and TBK1-dependent signalling pathway in chicken macrophages and evidencing the cross-species utility of these two pharmacological inhibitors (Figure 3D). As with DNA stimulation, there was no measurable impact of cGAMP treatment on phagocytosis in HD11 cells (Figure 3E).

**Figure 3.**
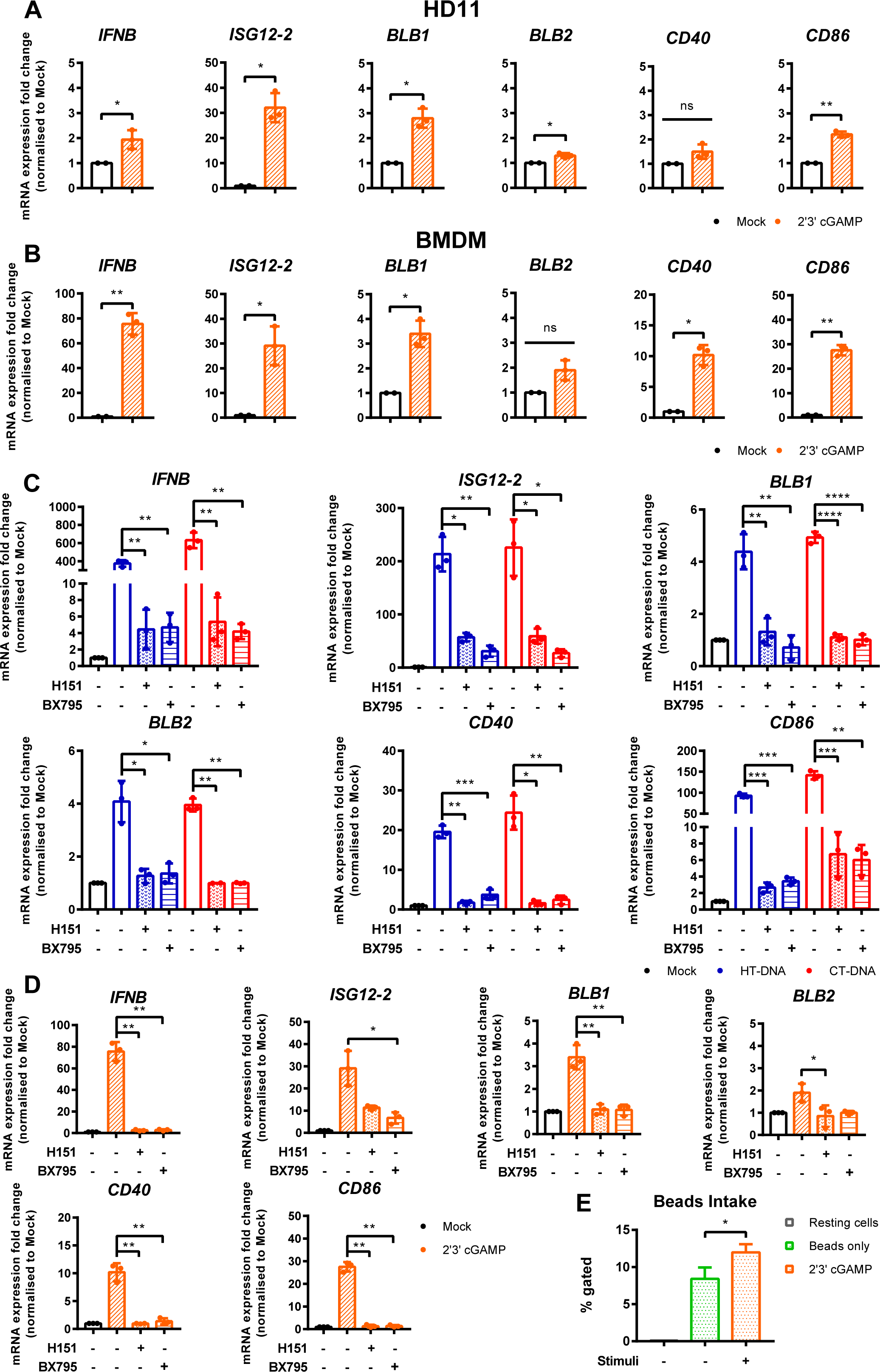
STING and TBK1 contribute to DNA-driven transcriptional responses in chicken BMDMs. (A) HD11 and (B) BMDM cells were treated with 2’3’cGAMP (10 μg/mL) and qRT-PCR carried out 6 h later for the indicated genes. (C) BMDM were treated with the STING inhibitor H-151 (10 uM) or TBK1 inhibitor BX795 (1 uM) for 1 h before transfection with HT-DNA and CT-DNA. 6 h later RNA was extracted and qRT-PCR carried out for the indicated genes. (D) BMDM were treated with the STING inhibitor H-151 (10 uM) or TBK1 inhibitor BX795 (1 uM) for 1 h before treatment with 2’3’cGAMP (10 μg/mL). 6 h later RNA was extracted and qRT-PCR carried out for the indicated genes. (E) HD11 cells were treated with 2’3’cGAMP (2.5 µg/mL) 6 h later phagocytosis was monitored by FITC-conjugated, zymosan coated bead uptake. *: p < 0.05, **: p < 0.01, ***: p < 0.001; ****: p < 0.0001; ns: no significant difference

### cGAS is essential for intracellular DNA-dependent IFN-I and MHC-II transcription in HD11 cells

To address the possibility that cGAS is a principle PRR responsible for sensing intracellular DNA in chicken macrophages, we generated HD11 knockout cell lines using CRISPR/Cas9 genome editing. To do this we analysed the annotated cGAS sequence in the current release of the *Gallus gallus* genome and designed gRNA sequences targeting regions of the gene which exhibited high conservation across multiple orthologues. By sequencing single cell clones we generated multiple cGAS knockout cell lines with two different gRNAs. By sequencing across the gRNA PAM target sites, we characterised indels to confirm the knockout status in these clones (eg Figure 4A). Stimulation of multiple cGAS knockout HD11 clones, each with a different indel, with DNA resulted in an abrogation of IFN-I and ISG transcription indicating that cGAS is a key PRR for sensing intracellular DNA in chicken macrophages (Figure 4B and Supplementary Figure 2). cGAS knockout also abrogated the upregulation of DNA-driven BLB1 stimulation, indicating the cGAS-dependent signalling is responsible for regulation of MHC class II transcription in this context (Figure 4B). These data were independent of IFNα pre-treatment, which enhanced IFN-I and BLB1 transcription in WT DNA-stimulated cells, but did not affect cGAS KO cells (Figure 4C). Consistent with the mammalian cGAS mechanism, stimulation of WT or cGAS KO cells with 2’3’-cGAMP resulted in robust IFN-I transcription, indicating IFN-I production by direct STING ligation was not affected by cGAS KO (Figure 4D). These data confirm the intracellular DNA PRR function of cGAS in chicken macrophages.

**Figure 4.**
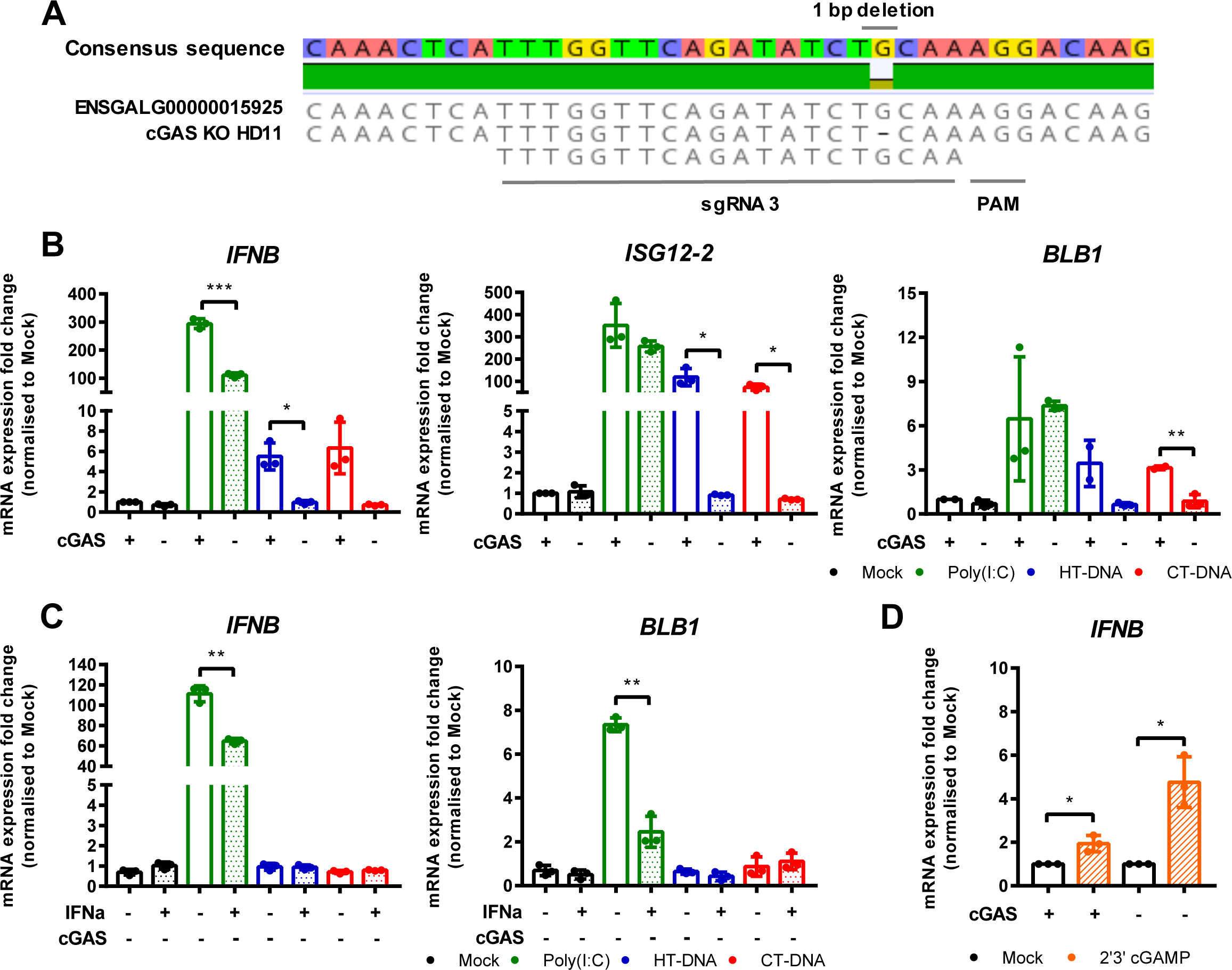
cGAS is essential for intracellular DNA-dependent IFN-I and MHC-II transcription in HD11 cells. (A) Example of identification of indel in clonally selected HD11 cGAS KO using NGS sequencing. (B, C) WT and cGAS KO HD11 cells were transfected with HT-DNA, CT-DNA (2 μg/mL) or Poly(I:C) (1 μg/mL) for 6 h and transcription of the indicated genes measured by qRT-PCR. (D) cGAS KO HD11 cells were primed with IFNα for 6h, transfected with HT-DNA, CT-DNA, or Poly(I:C) and transcription of IFNB and ISG12.2 measured by qRT-PCR 6 h later (E) WT or cGAS KO cells were treated with 2’3’cGAMP (10 μg/mL) and transcription of IFNB measured by qRT-PCR 6 h later. *: p < 0.05, **: p < 0.01, ***: p < 0.001; ****: p < 0.0001; ns: no significant difference

### STING is essential for intracellular DNA-dependent IFN-I transcription in HD11 cells

In parallel, using the same methodology, we generated multiple STING knockout HD11 cell lines (Figure 5A). Stimulation of these cells with DNA phenocopied the cGAS knockout lines, confirming the function of chicken STING downstream of cGAS in the intracellular DNA sensing pathway (Figure 5B, Supplementary Figure 3). These data are consistent with the presence of a cGAS/STING pathway in HD11 cells and, in concert with the data using H151 in BMDMs, indicate the function of STING as a critical adaptor protein for intracellular DNA sensing in chicken macrophages.

**Figure 5.**
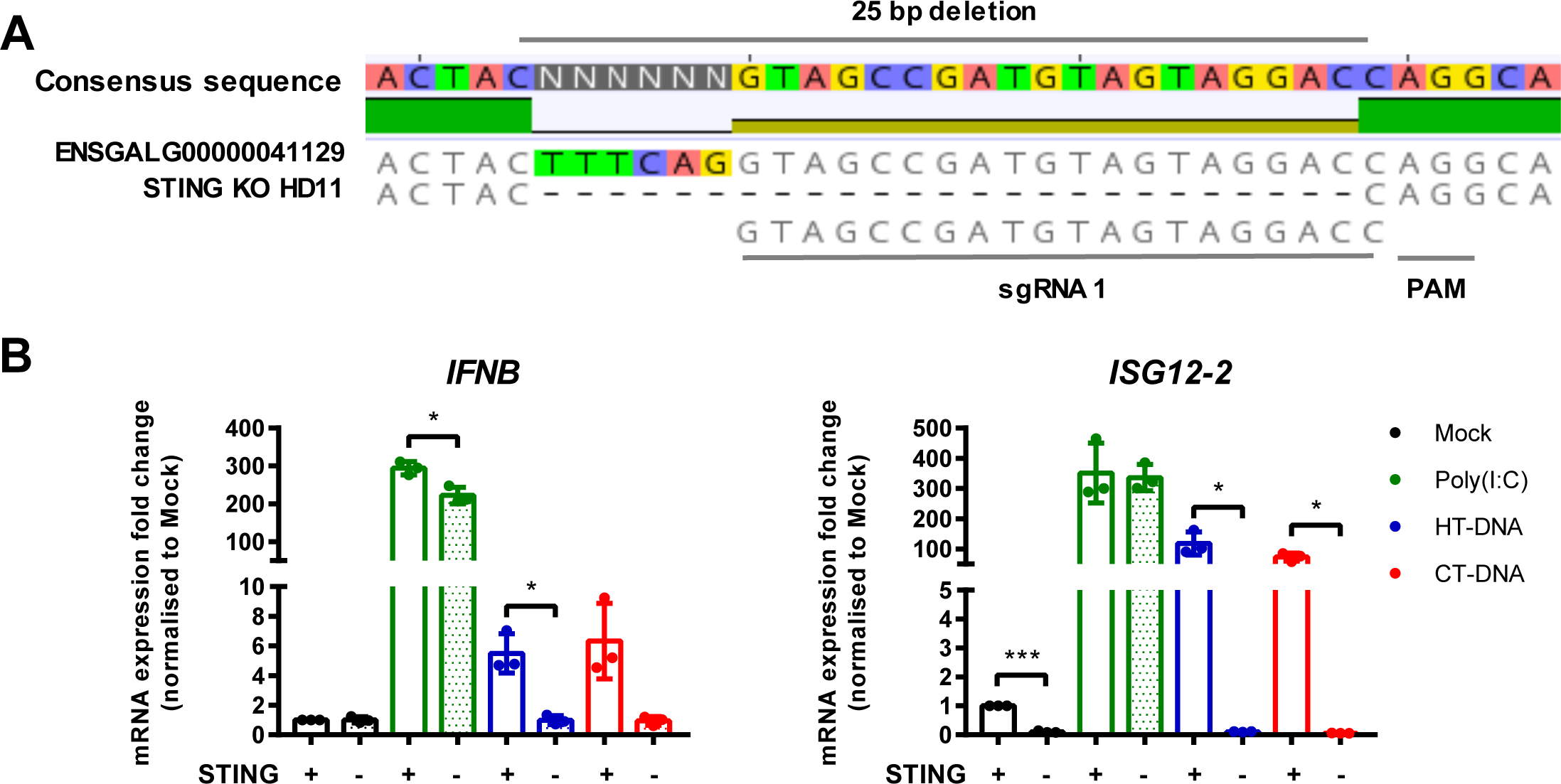
STING is essential for intracellular DNA-dependent IFN-I transcription in HD11 cells. (A) Example of identification of indel in clonally selected HD11 STING KO using NGS sequencing. (B) WT and STING KO HD11 cells were transfected with HT-DNA, CT-DNA (2 μg/mL) or Poly(I:C) (1 μg/mL) for 6 h and transcription of the indicated genes measured by qRT-PCR 6 h later. *: p < 0.05, **: p < 0.01, ***: p < 0.001; ****: p < 0.0001; ns: no significant difference

### Fowlpox triggers a cGAS / STING dependent DNA sensing pathway in HD11 cells

FWPV replication exposes large quantities of DNA to the cytoplasm of infected cells making it a prime target for intracellular DNA sensing PRRs. Despite this, using the wild-type vaccine strain FP9 we, and others (Giotis and Skinner, 2019; Laidlaw et al., 2013), observe little or no IFN-I transcription in infected cells, and indeed a downregulation of IFN and MHC transcription (Figure 6A). The lack of IFN-I response in poxvirus infected cells is likely due to the presence of numerous virally-encoded suppressors of PRR signalling and IFN-I production (Laidlaw et al., 2013; Smith et al., 2013), hence deletion of specific innate immunomodulators from the viral genome can result in a virus that stimulates host IFN-I signalling. We made use of FWPV mutants FPV012 and FPV184 ((Laidlaw et al., 2013, Giotis & Skinner, unpublished), each deficient in single genes that are proposed immunomodulators, and both of which induce IFN-I production from infected cells (Laidlaw et al., 2013), including HD11 cells (Figure 6B). In the absence of cGAS or STING the transcription of IFN-I, ISG12.2, BLB1 and CD40 by FPV184 or FPV012 was significantly lower at 24h post infection (Figure 6B), despite robust infection of HD11 cells by all three virus strains (Figure 6C), indicating that FWPV is sensed in infected cells by the DNA sensing PRR cGAS and that the cGAS/STING pathway is responsible for FWPV-induced IFN-I production and MHC-II transcription.

**Figure 6.**
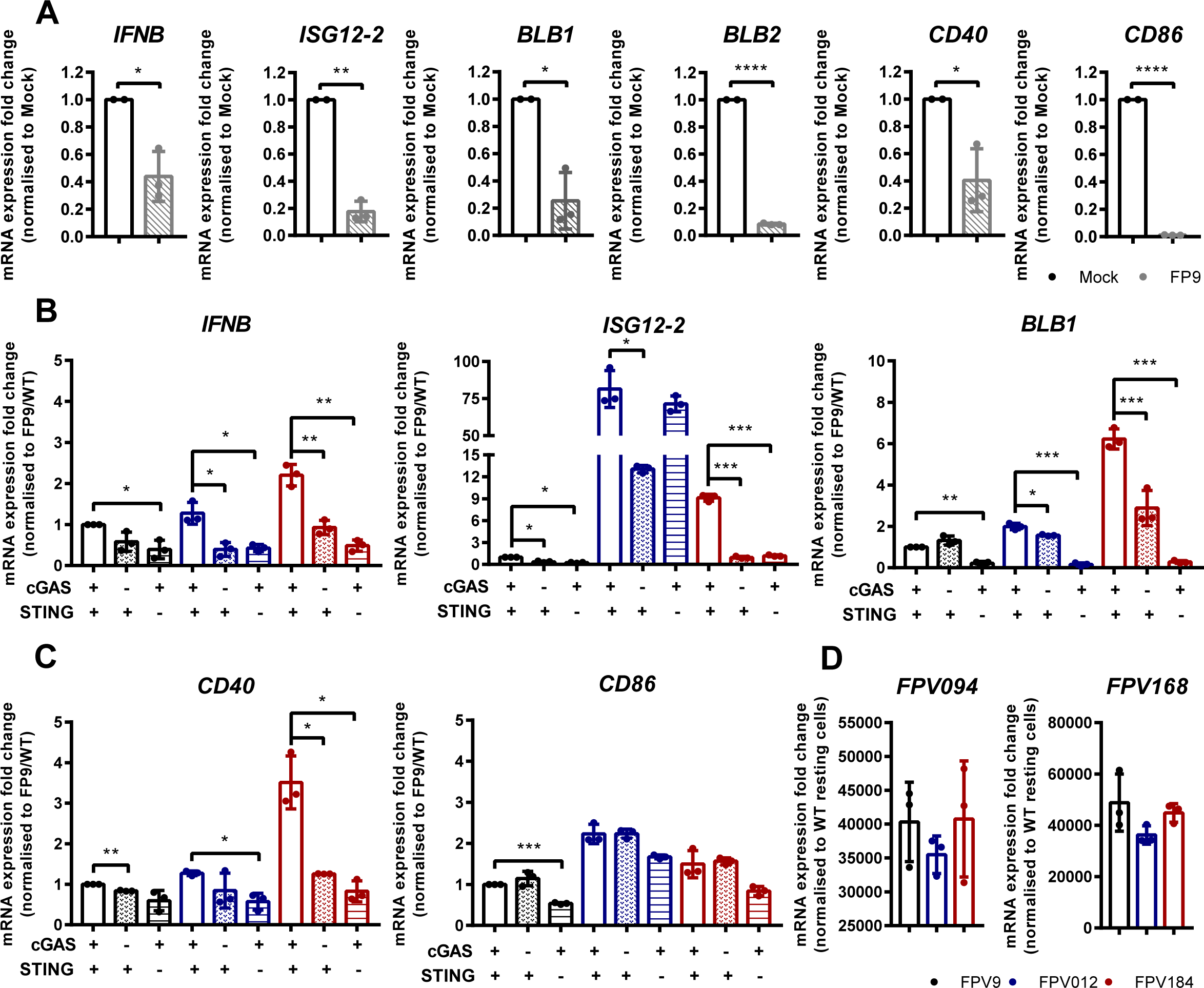
Fowlpox triggers a cGAS / STING dependent DNA sensing pathway in HD11 cells. (A) HD11 cells were infected with FWPV strain FP9 at a multiplicity of infection of three. 24 h later RNA was extracted and qRT-PCR carried out for the indicated genes. (B, C) HD11 WT, cGAS or STING KO cells were infected with FP9, FPV012 or FPV184 at a multiplicity of infection of three. 24 h later RNA was extracted and qRT-PCR carried out for the indicated genes. *: p < 0.05, **: p < 0.01, ***: p < 0.001; ****: p < 0.0001; ns: no significant difference

## Discussion

The ability of innate immune cells to detect virus infection is dependent on a set of PRRs that directly bind viral nucleic acids. Macrophages act in this context as tissue-resident sentinel sensors of infection that express a broad repertoire of PRRs and mount a rapid and robust innate immune response to viruses and other pathogens. Indeed intracellular DNA sensing was first described in macrophages (Stetson and Medzhitov, 2006). As well as interferon and cytokine production, activated macrophages use effector functions for pathogen clearance and for activation of adaptive immunity. In mammalian systems the signalling outputs downstream of intracellular DNA detection in macrophages include IRF-dependent IFN and cytokine production and cell death driven by the AIM2 inflammasome. In chicken macrophages, which lack AIM2, we find that intracellular DNA sensing produces IFN but doesn’t result in measurable cell death, rather it upregulates a specific set of antigen presentation machinery including the MHC-II gene BLB1 and co-stimulatory molecules, providing a direct link between anti-viral innate sensing and the initiation of adaptive immunity.

During DNA virus infection, the cGAS/STING-dependent signalling pathway is triggered by viral DNA, resulting in type-I interferon production via activation of TBK1 and the IRF family of transcription factors. Although well defined in mammalian systems, the function of chicken cGAS and STING has only more recently been identified (Gao et al., 2018; Vitak et al., 2016). FWPV is an avian poxvirus that causes skin lesions and respiratory infections and can infect multiple cell types including macrophages (Williams et al., 2010). Here we show that the cGAS/STING pathway in chicken macrophages can sense FWPV infection and is responsible for the IFN-I response as well as for upregulation of BLB1.

In order to escape detection and evade host anti-viral responses, poxviruses like FWPV encode a broad range of immunomodulatory proteins that target PRR signalling pathways resulting in these viruses being able to effectively inhibit IFN production from infected cells. These immune evasion mechanisms mask the signalling outputs of PRR signalling during infection with wild type poxviruses. To overcome this issue, we used two mutant FWPVs with deletions in individual genes that block IFN-I production during infection. Infection of cells with FPV184 and FPV012 (Giotis et al., 2016) resulted in interferon and ISG transcription, which was lost in cGAS and STING knockout lines. FWPV DNA is therefore sensed by the cGAS/STING pathway and the downstream signalling response leading to IFN-I production is effectively blocked by the wild type virus.

Birds occupy the same habitats as mammals, have comparable ranges of life span and body mass, and confront similar pathogen challenges, yet birds have a different repertoire of organs, cells, molecules and genes of the immune system compared to mammals (Kaiser, 2010). It is increasingly evident that the immune system of avian species is rather different from those of model mammalian species. Untested extrapolation from mammalian systems cannot provide the quality of knowledge that is required for understanding host-pathogen relationships in birds. Here we find that the signalling downstream of chicken cGAS leading to IFN-I transcription is similar to that found in mammalian systems. The presence of orthologues of STING and TBK1 in the chicken genome and their functional inhibition by small molecule compounds (H151 and BX795) is indicative of mechanistic signalling pathway conservation. The chicken genome also contains an orthologue of IRF3, which is the main transcription factor downstream of STING/TBK1 activation, although chicken IRF7 (as this gene is annotated) is not equivalent to mammalian IRF3 or IRF7 and may be considered more as a hybrid these two genes (Grant et al., 1995). It is likely that chicken IRF7 and TBK1 are recruited by STING following 2’3’-cGAMP ligation and that subsequent phosphorylation, dimerisation and nuclear translocation of IRF7 leads to DNA-induced IFN-I transcription (Cheng et al., 2019; Gao et al., 2018). Recent evidence has implicated chicken cGAS and STING in avian antiviral defence, in particular against Marek’s Disease Virus (MDV) and chicken adenovirus 4 (Li et al., 2019; Wang et al., 2020) in fibroblasts. Using CRISPR/Cas9 technology to knockout STING and cGAS in a transformed monocytic cell line (HD11) and complementing these data in primary macrophages with pharmacological inhibitors we have been able to show this cGAS/STING/TBK1 pathway is active in chicken macrophages. The use of primary cells in this context is important as transformation or immortalisation can significantly alter PRR pathways so as to obscure physiological signalling mechanisms.

IFN-I is one of the most effective anti-viral innate immune mediators. Secretion and subsequent ISG transcription induced by autocrine and paracrine IFN receptor signalling sets an anti-viral/inflammatory state in infected and bystander cells. As an example, chicken IFNβ was shown to be an autocrine/paracrine pro-inflammatory mediator in chicken macrophages (Garrido et al., 2018), with direct effects in macrophage effector functions. Nucleic acid sensing PRRs therefore provide a rapid and potent innate response helping to combat infection and reduce viral spread in infected tissues. At the same time, innate immune responses can initiate and amplify adaptive immune responses for example, by regulating functions of antigen presenting cells (APCs), promoting cross-priming and stimulating antibody production (Desmet and Ishii, 2012; Loré et al., 2003; Schulz et al., 2005). In both mammals and birds, macrophages are key regulators of adaptive immunity as principle APCs. By processing and presenting antigen to T and B cells, macrophages directly trigger adaptive responses. The discovery that cGAS/STING signalling can directly regulate the transcription of MHC genes in macrophages provides further evidence linking PRR signalling with the activation of adaptive immunity during infection. It remains to be explored exactly how the transcription of BLB1 and BLB2 is regulated by cGAS/STING signalling. In tissues, macrophages survey the local environment for infection and damage. In this context, macrophage effector functions may be modulated by the presence of innate immune mediators in the tissue. The priming effect of IFNα as an enhancer of macrophage DNA sensing, by upregulating STING expression, suggests a possible mechanism of bystander surveillance. Tissue resident macrophages may respond to signals, including IFN-I and cGAMP, secreted from virally infected stromal cells by enhancing specific effector functions appropriate to defend against viral infection in the tissue (Ablasser et al., 2013c; Schadt et al., 2019).

Our data adds to the list of chicken cGAS/STING functions in sensing of avian DNA viruses such as MDV and Adenovirus 4 that replicate in the nucleus or FWPV that replicates in the cytoplasm, and in the regulation of macrophage effector functions. The ability of this pathway to sense a broad range of DNA viruses that replicate in different compartments in avian innate immune cells indicates that this pathway is a primary DNA sensing mechanism for DNA viruses in chickens.

---

**Supplementary Figure 1.**
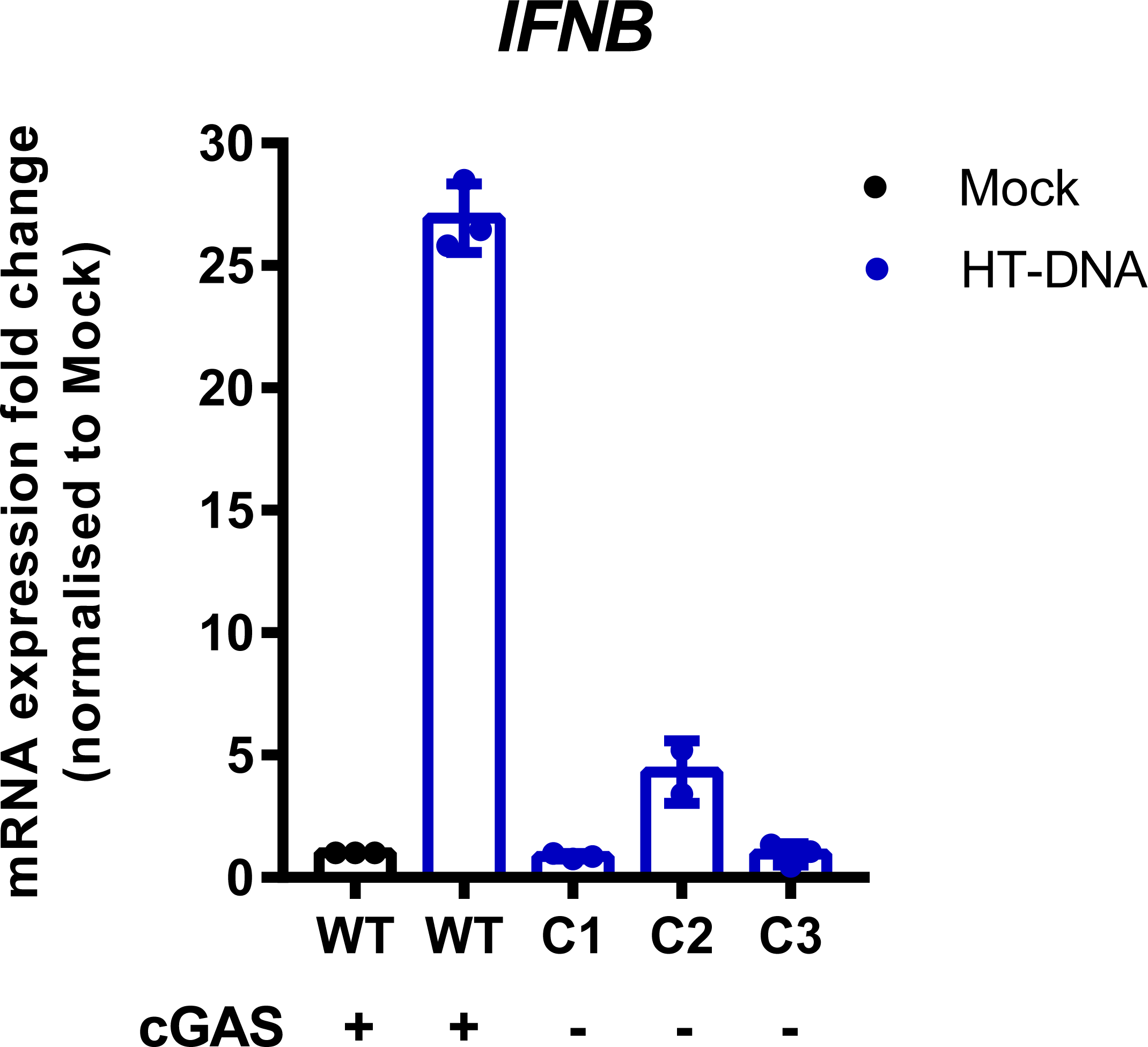
Effect of IFNα priming on expression levels of STING and IRF7 in BMDM and HD11. BMDM or HD11 cells were treated with IFNα for 6 h and transcription of STING and IRF7 measured by qRT-PCR 6 h later.

**Supplementary Figure 2.**
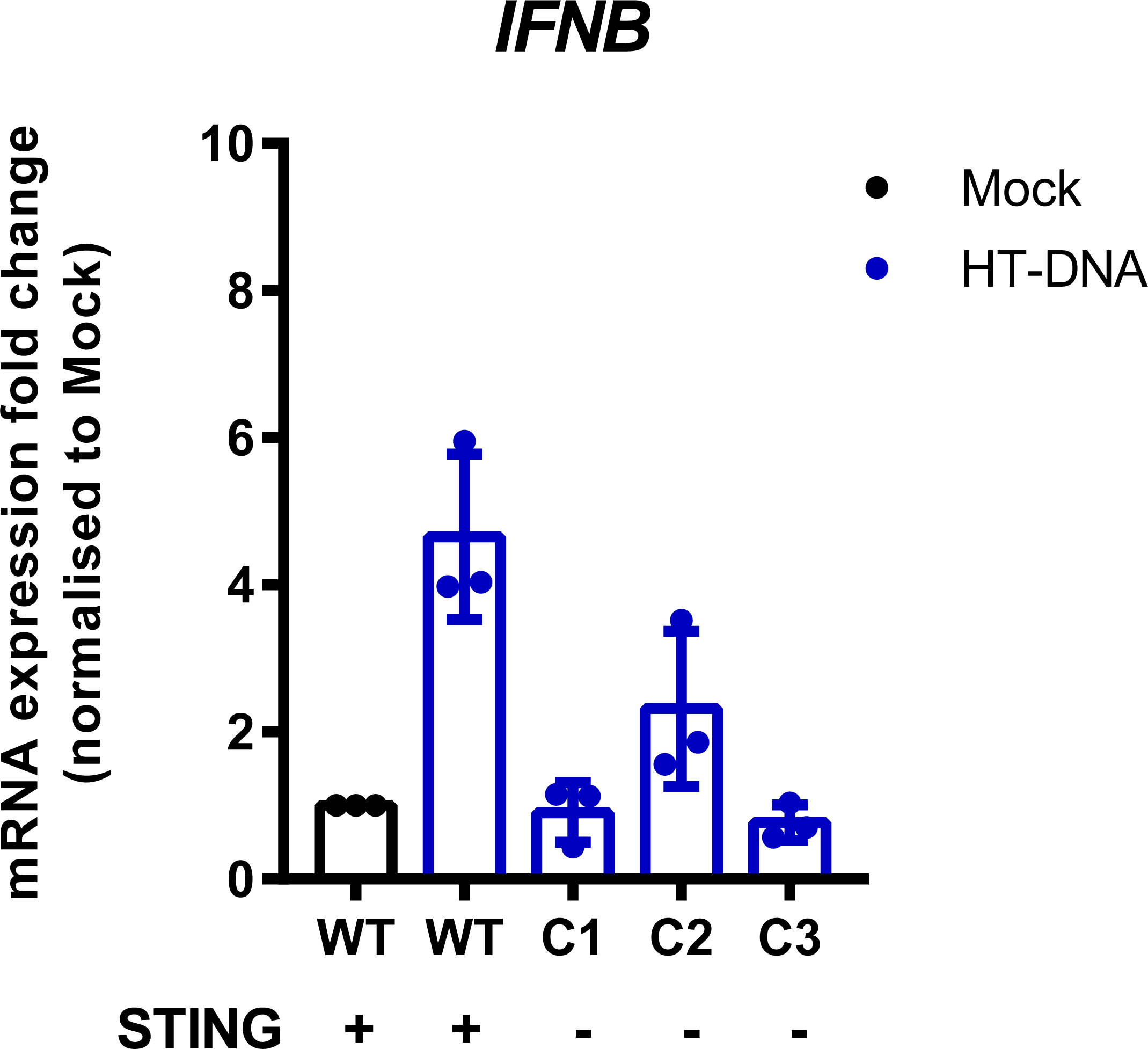
cGAS is essential for intracellular DNA-dependent IFN-I transcription in HD11 cells. WT or three individual cGAS knockout clones with different indels were stimulated with HT-DNA (2 μg/mL) and IFNB transcription measured by qRT-PCR 6 h later.

**Supplementary Figure 3.**
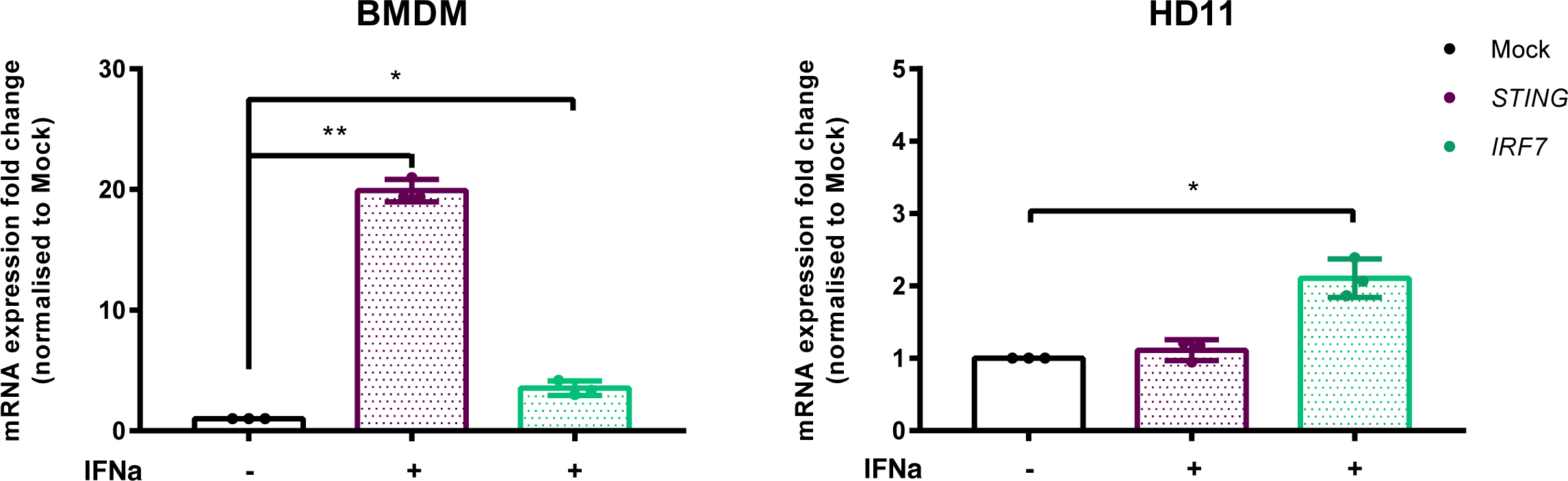
STING is essential for intracellular DNA-dependent IFN-I transcription in HD11 cells. (A) WT or three individual cGAS knockout clones with different indels were stimulated with HT-DNA (2 μg/mL) and IFNB transcription measured by qRT-PCR 6 h later.

## Author Contributions

BF and CB provided the funding and supervised the work. MO, DR, RG, VG and EK performed the experiments and statistical analysis. SG and MS generated the mutant fowlpox viruses. BF, CB, RG designed the study and wrote the manuscript.

## Acknowledgments

This manuscript has been released as a Pre-print at BioRxiv. We thank the experimental facility PFIE (Plateforme d’Infectiologie Expérimentale, Centre INRAE Val de Loire, Nouzilly, France) for providing the animals used for the isolation of primary macrophages.

